# AAV-Mediated *In Vivo* CAR Gene Therapy for Targeting Human T Cell Leukemia

**DOI:** 10.1101/2021.02.15.431201

**Authors:** Waqas Nawaz, Bilian Huang, Shijie Xu, Yanlei Li, Linjing Zhu, Zhiwei Wu, Xilin Wu

## Abstract

Chimeric antigen receptor (CAR) T cell therapy is the most active field in immuno-oncology and brings substantial benefit to patients with B cell malignancies. However, the complex procedure for CAR T cell generation hampers its widespread applications. Here, we describe a novel approach in which human CAR T cells can be generated within the host upon injecting an Adeno-associated virus (AAV)vector carrying the CAR gene, which we call AAV delivering CAR gene therapy (ACG). Upon single infusion into a humanized NCG tumor mouse model of human T cell leukemia, AAV generates sufficient numbers of potent *in vivo* CAR cells, resulting in tumor regression; these in vivo generated CAR cells produce antitumor immunological characteristics. This instantaneous generation of *in vivo* CAR T cells may bypass the need for patient lymphodepletion, as well as the *ex vivo* processes of traditional CAR T cell production, which may make CAR therapy simpler and less expensive. It may allow the development of intricate, individualized treatments in the form of on-demand and diverse therapies.

**Significance Statement:** AAV can generate enough CAR cells within the host. That act as a living drug, distributed throughout the body, and persist for weeks, with the ability to recognize and destroy tumor cells.

## Introduction

Chimeric antigen receptor (CAR) T cells have achieved significant clinical recognition with remarkable responses documented for autologous CD19 (a B cell-specific antigen) redirected T cells for the treatment of patients with relapsed or refractory B cell malignancies [1]. In 2017, the US Food and Drug Administration (FDA) approved two autologous CD19-CAR T cell products to treat certain types of relapsed or refractory large B cell lymphoma and acute lymphoblastic leukemia [2]. However, CAR T cell therapy still faces skepticism and has plenty of room for improvement, including challenges ranging from understanding the mechanisms and management of innate and acquired resistance and cytokine release syndrome (CRS) to neurologic toxicity and establishment of more efficient, less expensive, and quicker mechanisms of manufacturing [3]. Current clinical-scale manufacturing of T lymphocytes involves complex processes of isolation, genetic modification, and selective expansion of redirected cells *ex vivo* prior to their infusion into patients. This makes the manufacturing process complicated, costly and associated with clinical hazards. Thus, the manufacturing of safe, affordable, easier and effective CAR T cells is critical for making this therapy widely accessible for a larger population [4]. Moreover, these complicated procedures are resource-intensive and require substantial technical expertise and can only be performed at a limited number of specialized centers worldwide [5]. The most crucial step to overcome these limitations is to create novel strategies for making this complicated and risky cell therapy affordable, safer and customizable. This may in turn enable CAR T cells to outcompete chemotherapy as front-line therapy.

Adeno-associated virus (AAV), a small, icosahedral nonenveloped virus of 25 nm in diameter containing a single-stranded DNA genome, has the potential to solve this problem. AAV has become an essential therapeutic gene delivery vector in recent years, it is frequently used to deliver genetic material into target cells to cure or treat disease, and is extensively used in clinical applications in academia and industry [6]. To date, almost 200 clinical trials using AAV have been completed or are ongoing [7].

To manipulate immune cells *in vivo*, the carrier needs to be taken up by cells and import the encoding DNA into the nucleus. Previous studies showed that AAV could transduce both dividing and non-dividing cells with superior infection efficiency, which enables AAV to be widely applied in gene therapy in vivo [8]. In our current study, we choose AAV-DJ (type2/type8/type9 chimera), which was developed by using DNA shuffling technology to produce hybrid capsids from eight distinct wild-type AAV [9]. Previously, we reported AAV-DJ to deliver antibody gene in vivo, where antibodies were persistent to express in cells for more than 9 weeks [10]. The capability to transduce different cell types is mainly determined by the AAV capsid proteins. AAV-DJ capsid was selected [9], which has higher tropism and extremely robust at transducing cells [9, 11].

We hypothesize that human CAR T cells can be generated to specifically kill target cells within the host upon injecting an AAV vector carrying the CAR gene, which we call AAV delivering CAR gene therapy (ACG). To prove this concept, we furnished an AAV vector with plasmid DNA encoding a CD4-CAR, which is composed of a single-chain antibody (scFv) specific for the extracellular domain of the CD4 antigen fused with CD28, 4-1BB, and CD3 zeta cytoplasmic signaling domains. We demonstrated that the modified AAV vector could safely deliver the CAR gene into the host cells of humanized NCG (NCG-HuPBL) mice and reprogrammed cells to express the CAR. As envisioned, the AAV vector generated enough CAR T cells *in vivo* and thus caused effective tumor regression and produced antitumor immunological characteristics. In addition, *ex vivo* experiments indicated that our AAV delivering the CD4-CAR potently eliminated tumor cells from adult T cell leukemia (ATL). The current approach may skip the technically challenging *ex vivo* procedures and enable the host to create its CAR T cells.

## Methods and materials

### Mice

NCG mice were purchased from GemPharmaTech. NCG mice were maintained in accordance with the Guide for the Care and Use of Laboratory Animals of the Medical School of Nanjing University. All experiments were performed according to the guidelines of the Institutional Animal Committee of Nanjing University.

### PBMCs and cell lines

Peripheral blood mononuclear cells (PBMCs) (derived from healthy donors visiting the Drum Tower Hospital, Nanjing University) were isolated from buffy coats by Ficoll-Hypaque gradient separation, and cultured in RPMI-1640, 10% heat-inactivated FBS and freshly added recombinant human interleukin-2. Excess PBMCs were cryopreserved until use. For stimulation, PBMCs were washed once in T cell medium without IL-2 and then suspended at a concentration of 10^6^/ml in T cell medium containing IL-2 (300 IU/ml) and anti-human CD3 antibody (1 μg/ml). Cells were incubated in 5% CO2 at 37°C for 2 days until AAV transduction.

AAV-293T (Thermo Fisher) and HEK-293T (ATCC) cells were maintained in Dulbecco’s modified Eagle medium (DMEM) containing 10% FBS and 2 mM glutamate. H929, MT2, and Jurkat cell lines (ATCC) were grown in RPMI-1640 with 10% heat-inactivated FBS. CD4 expression was authenticated by flow cytometry in MT2 and Jurkat cell lines prior to experiments. All cell culture media contained 100 U/ml penicillin-streptomycin (Gibco, Invitrogen). All cell lines were maintained in an environment of 37°C and 5% CO_2_.

### Construction of AAV-CAR vectors

A recombinant AAV-2 ITR-containing plasmid (pAAV-MCS) with an expression cassette containing inverted terminal repeats was used for AAV generation. To generate AAV-CD20CAR and AAV-CD4CAR, we used the sequence of CD20-binding scFv from NCBI (GenBank, KM043780) and the sequence of Hu5A8 specific for the CD4 backbone with a 20-mer (Gly4Ser) adopted from our previous study [10]. The scFv was fused with CD28 hinge-transmembrane regions linked to 4-1BB (CD137) intracellular domains and the CD3 zeta intracellular domain [12]. CAR DNA (Hu5A8-CD28-41BB-CD3 or Rit-CD28-41BB-CD3) was cloned into the ITR/MCS-containing vector. Plasmid sequence and construction were confirmed by sequence analysis and restriction enzymes. Stocks of pAAV-Hu5A8-CD28-41BB-CD3 and pAAV-Rit-CD28-41BB-CD3 were prepared as pAAV vectors of interest (VOIs).

### AAV production

AAV of DJ subtype was prepared by polyethyleneimine (PEI)-based triple transfections of pAAV-VOI, pHelper (helper plasmid) and pAAV-DJ (catalog number VPK-430-DJ, Cell Biolabs), as described [13]. Briefly, the medium of 80–90% confluent AAV-293T cells in 15 cm plates was changed to fresh DMEM 2 hours before transfection. Plasmids at a ratio of 1:1:1 (25 µg each plasmid for a 15 cm plate) were mixed with 2 ml Opti-MEM in a 15 ml tube. Three hundred microliters of PEI (Polysciences Inc.) (1 mg/ml) was added to 2 ml Opti-MEM in another 15 ml tube and was mixed by vortexing and incubated for 15 min at room temperature, after which the mixture was gently added to the serum-free medium in the plate and incubated in a standard 5% CO2 and 37°C incubator. The supernatant was collected 24, 48, 72, and 96 hours posttransfection, and the debris was cleared by centrifugation and filtration (0.45 µM). AAV was concentrated by centrifuging (1000g for 5min) with 10% polyethylene glycol 8000 and 14% 4 M NaCl (by volume) and incubated overnight at 4°C. The next morning, the sample was transferred to 50 ml conical tubes and centrifuged at 4000 g for 30 min at 4°C. The AAV pellet was resuspended in PBS and stored at −80°C. We used qPCR to determine the number of genome-containing particles in an AAV prep using SYBR green technology, adopted from our previously reported protocol [10]. The detailed procedure of determining the genome-containing particles in an AAV prep using SYBR green technology is explained in the supplementary information.

### AAV transduction of HEK-293T and T cells

Three different doses of AAVs were administered: 1×10^4^ AAVs/cell was considered AAV low (L), 1.5×10^4^ AAVs/cell was considered AAV middle (M), and 1×10^5^ AAVs/cell was considered AAV high (H). The day before transduction, HEK-293T cells were trypsinized and counted, and 1-4 × 10^5^ cells were plated in 2.0-3.0 ml complete culture medium and incubated at 37°C overnight. Sixteen hours later, purified AAV (AAV-CD4CAR or AAV-CD20CAR) was added to each well at different doses by gentle swirling/mixing. Twenty-four hours post-transduction, fresh media containing 20% FBS was added, and the cells were cultured for 2-3 days and then analyzed by FACS, or fluorescence microscopy.

The T cells were activated and infected with AAV-CAR as described [14]. Briefly, 10^6^ cells were plated on day 0 and activated for two days in the presence of 300 IU/ml IL-2 and 1 μg/ml anti-human CD3 antibody. On day 2, purified AAV (AAV-CD4CAR or AAV-CD20CAR) was added to each well at different doses by gentle swirling/mixing and incubated at 37°C in the presence of 5% CO2. The next morning following transduction, fresh media containing 20% FBS was added, and the cells were cultured for 2-3 days and then analyzed by FACS or florescence microscopy.

### Flow cytometry

All antibodies for flow cytometry were purchased from BioLegend or Invitrogen. Cells were stained with target antibodies (according to the experiment’s requirement), whereas for CAR expression, target antigen (CD4Fc protein) was used. After 30 minutes at 4°C, cells were washed once, suspended in FACs buffer, and stained with Alexa flour488 anti-rabbit for 30 minutes at 4°C. For in vivo samples, cells were lysed by RBC lysis buffer (BD), by incubating the cells with 1X lysis buffer at room temperature for 10 minutes. Labeled cells were washed twice and suspended in FACs buffer. All cells were measured and sorted by using a NovoCyte FACS, and analyzed with FlowJo software version 10.2.

### CAR T cell-mediated cytotoxicity assay using CFSE and 7-AAD

Cytotoxicity assays were carried out using CFSE and 7-AAD, as previously described [15]. Briefly, CFSE-labeled target cells were incubated with different doses of AAV-CD4CAR- and AAV-CD20CAR-infected PBMCs for 24 hrs at 37°C. After 24 hours of incubation, 7-amino-actinomycin D (7-AAD; BD Pharmingen) was added, as recommended by the manufacturer. The fluorescence was analyzed by flow cytometry. Target cell cytotoxicity was calculated using the following formula: cytotoxicity = 100 x [(CFSE-labeled dead target cells)/ (CFSE-labeled dead target cells + CFSE-labeled lived target cells)].

### *In vivo* NCG-HuPBL mouse model for the *in vivo* CD3^+^CD4^+^ depletion assay

NCG mice were purchased from GemPharmaTech. NCG mice were maintained in accordance with the Guide for the Care and Use of Laboratory Animals of the Medical School of Nanjing University. All experiments were performed according to the guidelines of the Institutional Animal Committee of Nanjing University. The humanized NCG (NCG-HuPBL) mouse model was developed as reported previously by our group [16]. Briefly, human PBMCs were obtained from healthy individuals’ peripheral venous blood and purified by Ficoll-Paque (Pharmacia, Piscataway, NJ) density gradient centrifugation as described above in detail. Freshly isolated PBMCs were resuspended in PBS, and 1.5×10^7^ PBMCs/mouse were injected intraperitoneally into NCG mice. PBMC engraftment was confirmed after 3 weeks by FACS, and mice with an appropriate percent ratio of CD3+CD4+ and CD3+CD8+ were considered successful NCG-HuPBL mouse models. A total of 5 × 10^6^ *ex vivo*-transduced AAV-CD4CAR T cells or 1×10^11^ AAV-CD4CAR T cells and 1×10^11^ AAV-CD20CAR T cells were injected into each Hu-NCG mouse according to the specified grouping. Mice were bled via the submandibular vein, and peripheral blood cell populations and CAR expression were analyzed periodically using flow cytometry.

### *In vivo* NCG-HuPBL tumor mouse model

After successfully engrafting PBMCs in NCG mice, we modeled T cell tumors by intraperitoneally injecting 2.5 × 10^6^ luciferase-expressing MT2 cells into each NCG-HuPBL mouse (generation of stable luciferase-expressing MT2 ATL tumor cell line is explained in Supplementary information). NCG-HuPBL mice injected with luciferase-expressing MT2 cells underwent *in vivo* bioluminescence imaging at various times as specified for each experiment. Luciferase-based bioluminescence imaging was performed with a NightOWL II LB 983 (Berthold) *in vivo* imaging system. Mice were first anesthetized by inhaling isoflurane and were maintained with 2% isoflurane during imaging procedures. Mice were injected (I.P.) with D-luciferin (prepared in DPBS) at 150 mg/kg and imaged immediately after anesthesia. Images were captured, and bioluminescence intensity and tumor area were quantitated using Indigo live imaging analysis (Berthold). Total flux and area were measured through the automated analysis method by Indigo (Berthold).

### Confocal microscopy

Samples from mice for confocal imaging were prepared as described previously [17]. Briefly, cells were stained for CD4-Fc protein and with an anti-CD45^+^ antibody for 1 hour. Next, the cells were washed and stained with anti-CD4-Fc secondary antibody for 1 hour. Following washing, the cells were transferred onto a fibronectin matrix (Sigma Aldrich) and fixed in 2% paraformaldehyde. Slides were mounted with #1.5 coverslips and Mowiol mounting medium and examined using an FV3000 confocal microscope.

### *In vivo* cytokine measurement

*In vivo* cytokines from the serum/plasma of mice were measured using the Cytometric Bead Array Human Th1/Th2 Cytokine Kit (BD), following the manufacturer’s instruction. The kit was used for the simultaneous detection of Human TNF, IL-2, IL-6, IL-4, and IL-10 in a single sample. The Cytometric Bead Array Human Th1/Th2 Cytokine Kit contains a mixture of various capture beads with different fluorescent intensities which are coated with capture antibodies specific for each cytokine. First, we mixed the beads coated with six specific capture antibodies. Next, 50 μL of the mixed captured beads, 50 μL of phycoerythrin (PE) detection reagent, and 50 μL of the unknown serum sample or standard dilutions were added sequentially to each assay tube and incubated for 2 h at room temperature in the dark. The samples were washed with 500 μL of wash buffer and centrifuged at 800g for 5 mint. The bead pellet was resuspended in 400 μL buffer after discarding the supernatant. All the samples were sorted on the NovoCyte FACS. Fluorescent intensities give the individual cytokine concentration. The standard curve for the CBA assay was determined using different dilutions. The data were analyzed with FCAP Array software provided by BD.

### Statistical analysis

Specific statistical tests and metrics (median, mean, standard error) used for comparisons, along with sample sizes, are described in the Results and Figure legends. All statistical analyses were performed using GraphPad Prism software version 6.0. ns, no significance; ****P < 0.0001; ***, P < 0.001; **, P < 0.01; *, P < 0.05.

## Results

### Design and characterization of AAV-CD4CAR

As schematically shown in Figure 1a, AAV-CD4CAR is the third-generation CAR construct composed of anti-CD4 scFv, hinge (H) and transmembrane regions, and CD28 and 4-1BB intracellular signaling domains in tandem with the CD3 zeta signaling domain. The CD4-CAR DNA molecule was subcloned into the pAAV-VOI plasmid, which was mixed with the plasmids pAAV-Helper and pAAV-DJ to generate AAV virus carrying the CD4-CAR gene (AAV-CD4CAR). To verify the AAV-CD4CAR construct, infected HEK-293T cells were subjected to Western blot and fluorescence-activated cell sorting (FACS) analysis. Immunoblotting with an anti-CD3 zeta monoclonal antibody showed bands of the expected size for the AAV-CD4CAR CD3 zeta fusion protein, while the control cells infected with AAV-GFP showed no bands (Fig. 1b). Furthermore, AAV-CD4CAR infection efficiency was analyzed through FACS using CD4-Fc protein (Fig. 1c). Moreover, AAV-CD4CAR infects cells with limited toxicity and high viability (Supplementary Fig. 1a-c).

**Figure 1.**
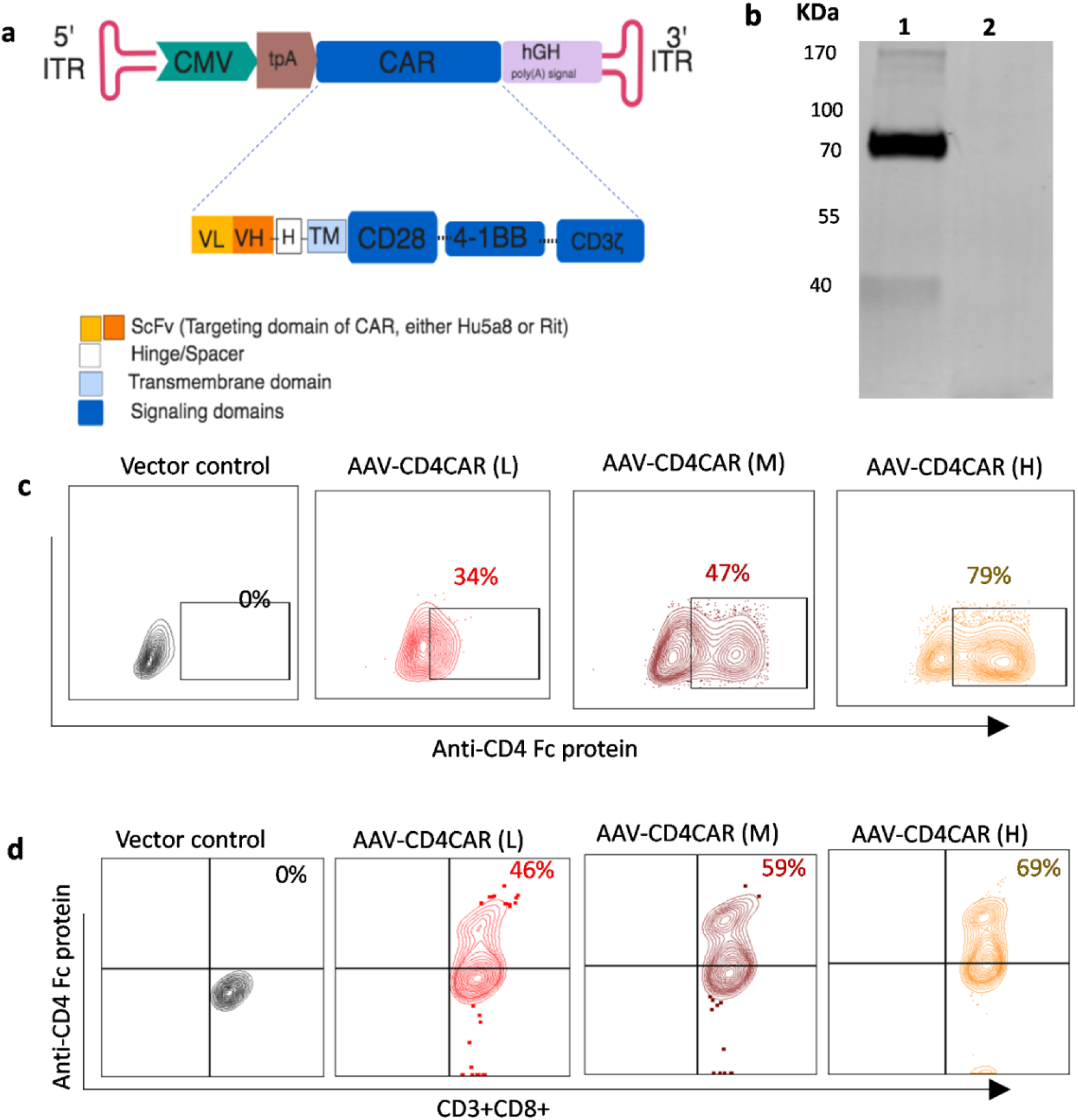
AAV-CD4CAR construct and characterization. **(a)** Schematic representation of the AAV encoding CD4CAR or CD20CAR. The third generation of the AAV-CAR contains a leader sequence, the anti-CD4 (Hu5a8) or the anti-CD20 scFv region (Rit), a hinge domain (H), a transmembrane domain (TM) and intracellular signaling domains from the following molecules: CD28 (coactivator), 4-1BB (coactivator), and CD3 zeta. **(b)** HEK293T cells were infected with AAV-CD4CAR (lane 1) and AAV-GFP (lane 2) for Western blot analysis, and at 48 hours post-transduction, they were probed with mouse anti-human CD3 zeta antibody, with the CD4-CAR band observed at the expected size (∼75 kDa) and a breakdown product observed at ∼37 kDa. However, no band was observed for AA V-GFP-transduced cells. **(c)** HEK293T cells were infected with different doses of AAV-CD4CAR. 48 hours post-infection, cells were stained with CD4-rFc protein followed by secondary antibody staining with Alexa Fluor 488-conjugated goat anti-rabbit IgG (H+L). **(d)** AAV-CD4 CAR expression on CD3+CD+8 cells. Activated PBMCs were infected with different doses of AAV-CD4CAR or uninfected (vector control). Forty-eight hours post-infection, cells were stained with CD4-rFc protein for CAR expression, followed by secondary antibody staining with Alexa-Fluor 488-conjugated goat anti-rabbit IgG (H+L). 1×10^4^ AAVs/cell was considered AAV low (L), 1.5×10^4^ AAVs/cell was considered AAV middle (M), and 1×10^5^ AAVs/cell was considered AAV high (H).

We characterized the effect of AAV-CD4CAR on T cells from healthy donors. Human T cells were activated with IL-2 and anti-human CD3 antibody and then infected with different doses of AAV-CD4CAR (MOI 1×10^4^, 0.5 × 10^5,^ and 1 × 10^5^). As expected, a dose-dependent expression of the CD4-CAR on T cells was observed (Figure 1d). We found that AAV-CD4CAR induced dose-dependent expression in three different healthy donor samples. However, the percentage of expression varied among donors, but the difference was not statistically significant (Supplementary Figure 2a). Upon more detailed phenotyping, we found that CD4+ T cells also expressed AAV-CD4CAR but at a significantly lower rate than CD8+ T cells (Supplementary Figure 2b).

In summary, these results confirmed that our AAV-CD4CAR contain the CD3 zeta on the intracellular end and the anti-CD4 scFv on the extracellular end, verifying that all other elements were present. Furthermore, AAV-CD4CAR infect the T cells and produce AAV-CD4CAR T cells in a dose-dependent manner.

### AAV-CD4CART cells are CD4 specific and eliminate leukemic cells *in vitro* and aggressive CD4^+^ ATL cells *ex vivo*

We furnished our AAV-CD4CAR with the scFv region of the Hu5A8 antibody specific for CD4, which we reported previously for HIV treatment [10]. We next evaluated the functional ability of AAV-CD4CAR to eliminate CD4^+^ cells in PBMCs from various donors (n=10) during T cell expansion following AAV-CD4CAR transduction. As expected, the groups treated with AAV-CD4CAR had a decreased CD3^+^CD4^+^ T cell subset within 2-3 days following AAV-CD4CAR transduction in a dose-dependent manner compared to the vector control group (Figure. 2b).

**Figure 2.**
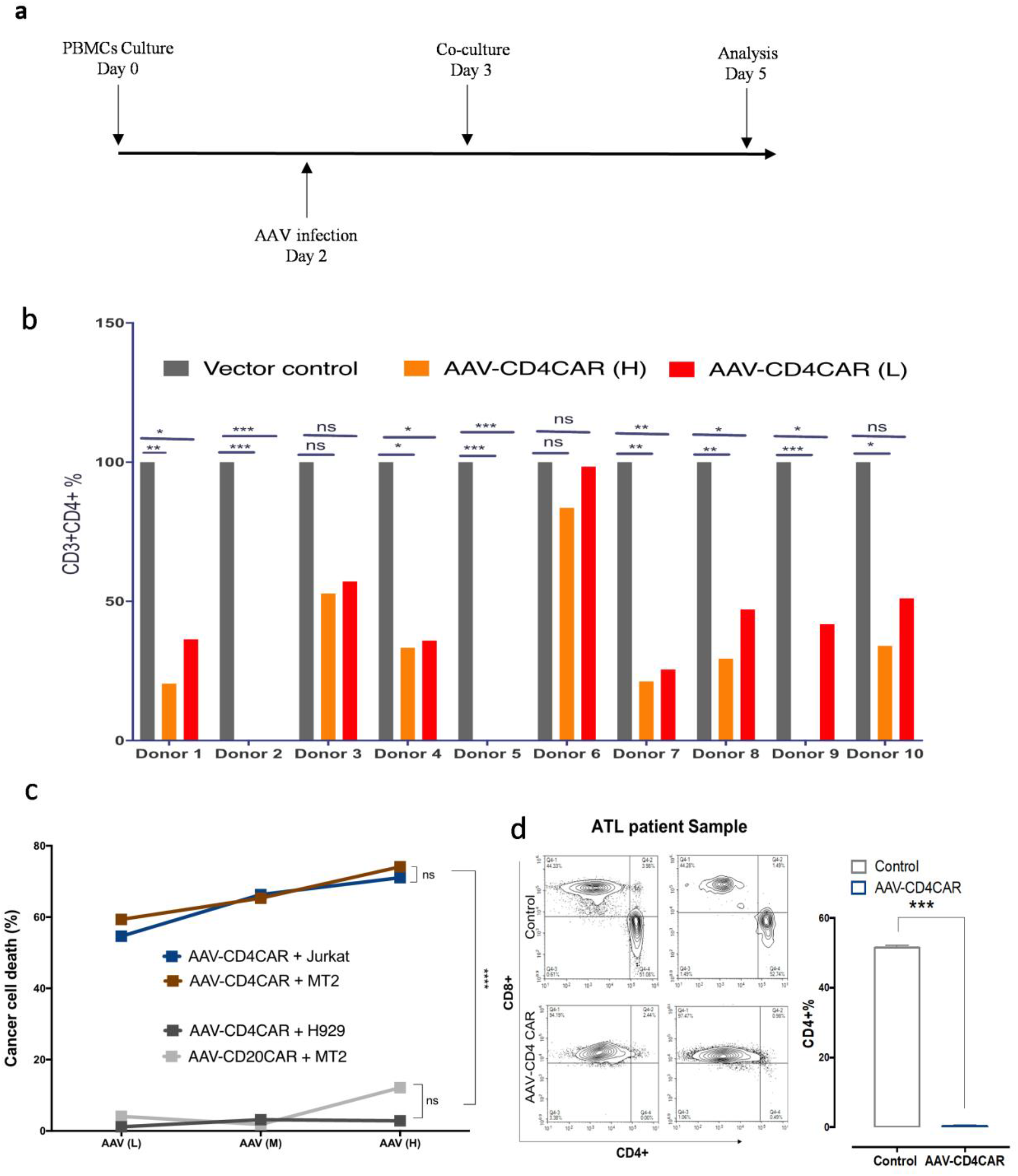
AAV-CD4CAR induces antitumor activity. **(a)** Schematic representation showing the *ex vivo* AAV-CAR production and co-culture assays. **(b)** AAV-CD4CAR depleted CD3+CD4+ cells in different donors (n=10) during the course of T cell expansion. Activated PB MCs from each donor were transduced with different doses of AAV-CD4CAR, and Vector control (untransduced). After 48 hours, the cells were analyzed by FACS using anti-human CD4, anti-human CD3, and anti-human CD8 antibodies. The CD3+CD4+ percentage was normalized to the vector control group (which represented 100%), and the CD3+CD4+ percentages of AAV-CD4CAR (H) and AAV-CD4CAR (L) was calculated by the percent difference from the vector control group. A two-way ANOVA multiple comparisons test within each don or was used for statistical analysis. ns, no significance; ***, P < 0.001; **, P < 0.01; *, P < 0.05. **(c)** AAV-CD4CAR T cells were co-cultured with CFSE-stained target cells, 48 hours post-co-culture, cells were stained with 7AAD and analyzed by FACS. Two-way ANOVA with the multiple-comparisons test was used to assess significance, ns, no significance; ****P < 0.0001. ***, P < 0.001; **, P < 0.01; *, P < 0.05. **(d)** ATL patient PBMCs were infected with AAV-CD4CAR (H) and with a control vector (samples were duplicated). 48 hours post-transduction, cells were analyzed by FACS for CD3+CD4+ depletion. An unpaired two-tailed t-test was used to calculate the significance of the difference. AAV-CD4CAR vs vector control, ***’P<0.0001.

AAV-CD4CAR was tested *in vitro* for its antileukemic activity using CD4-positive MT2 and Jurkat cells as targets. In addition, the CD4-negative cell lines H929 and AAV-CD20CAR served as a negative control. The MT2 cell line is an adult T cell leukemia (ATL) cell line with aberrant CD4 expression established by co-culture of normal human cord leukocytes with leukemic cells from ATL patients [18]. Jurkat cells are immortalized T lymphocytes obtained from a patient with T cell leukemia [19]. CD4 expression on MT2 and Jurkat cells was counterchecked by flow cytometry prior to experiments. H929 is a B cell line with CD38 expression established from a malignant effusion in a patient with myeloma [20]. 48 hours after co-culture with human PBMCs infected with various doses of AAV-CD4CAR, MT2, and Jurkat cells were effectively lysed in a dose-dependent manner. However, neither the CD4^-^ cell line (H929) nor the MT2 cells treated with a CD20-CAR were lysed (Figure. 2c). Next, we performed an *ex vivo* experiment to evaluate the ability of AAV-CD4CAR to kill primary ATL cells. CD3^+^CD4^+^ primary cells from an unclassified and chemotherapy-resistant ATL patient were isolated *ex vivo* and served as a target population. CD3^+^CD4^+^ ATL cells were completely lysed (depleted to ∼ 0%) by AAV-CD4CAR 48 hours post-infection, while AAV-CD20CAR was unable to lyse CD3^+^CD4^+^ ATL cells (Figure. 2d). In summary, these results show that AAV-CD4CAR-infected cells exhibited dose-dependent, profound, and specific antitumor activity against T cell leukemia cell lines. Similarly, they could also completely lyse the tumorigenic CD3^+^CD4^+^ population in samples from ATL patients in *ex vivo* coculture. Altogether, these results signify that AAV-CD4CAR-infected cells are specific for CD4^+^ cells.

### CAR T cells generated from in vivo reprogramming of lymphocytes via AAV and their antitumor immunological characteristics

We hypothesized that our AAV carrying CAR gene construct could reprogram lymphocytes and are able to generate *in vivo* CAR T cells to kill target cells in a process that we called ACG. To experimentally prove our ACG concept, we transplanted human PBMCs into NCG mice, and after successful engraftment of PBMCs, we injected 1 × 10^11^ AAV-CD4CAR into NCG-HuPBL mice. Interestingly, FACS analysis of blood collected after 48 hours showed that the AAVs started infecting the immune cells, and 9% of CD45^+^ cells expressed the CD4-CAR (Figure. 3a). We further confirmed CD4-CAR expression on CD45^+^ cells through confocal microscopy. Infused AAV-CD4CAR were taken up by lymphocytes, and the CD4-CAR could be observed on the lymphocyte membrane (Figure. 3a).

**Figure 3.**
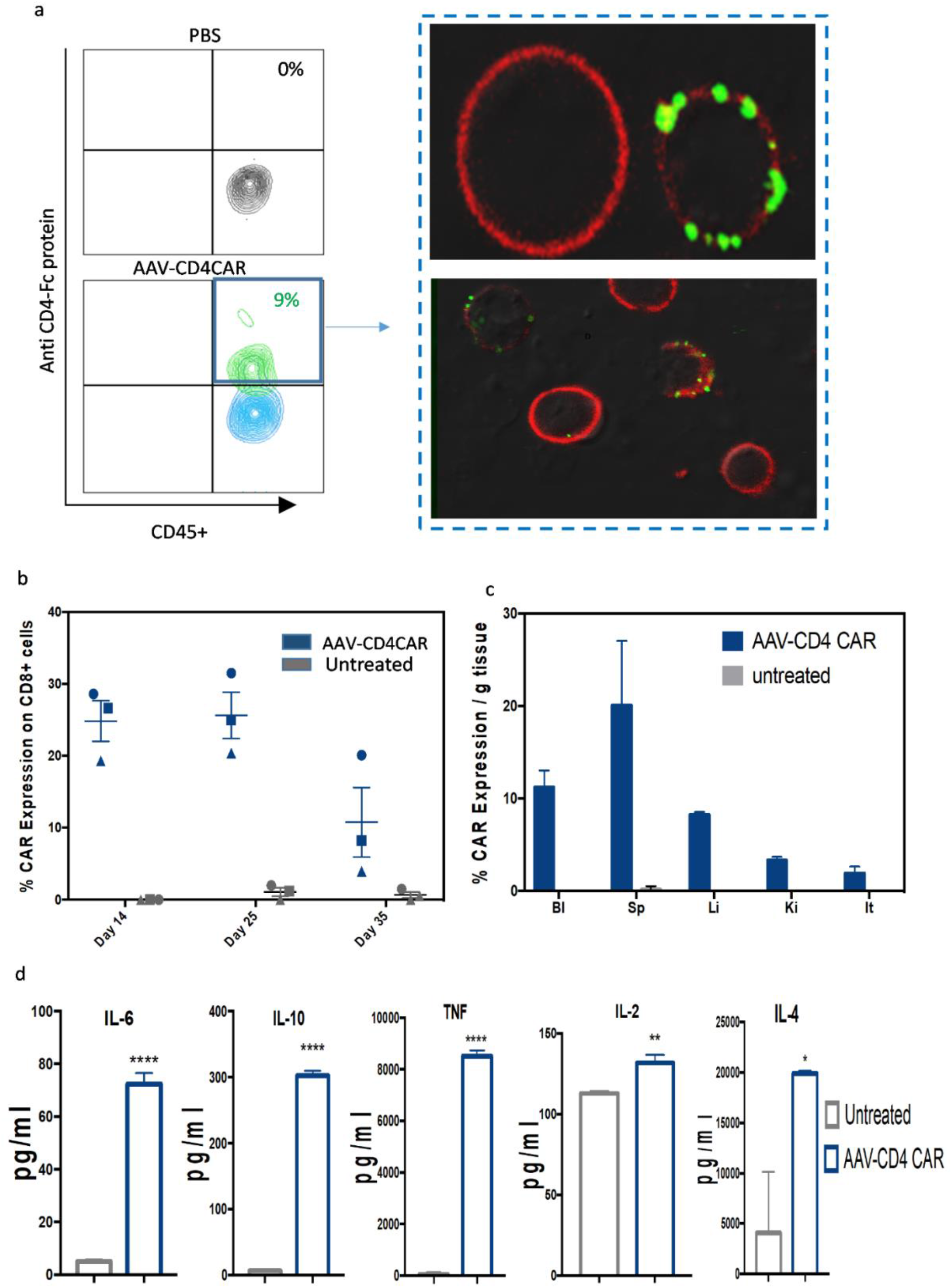
AAV-CD4CAR reprograms immune cells and generates *in vivo* tumor-specific AAV-CD4CAR T cells. **(a)** Representative flow cytometry results showing the expression of AAV-CD4CAR on CD45+ cells 48 hours after infusing a single dose of 1 × 10^11^ AAV-CD4CAR into NCG-HuPBL mice (n=3). The right panel shows the confocal imaging of CD45+ cells expressing AAV-CD4CAR expression. The red shows the cellular membrane, whereas the green dots represent AAV-CD4CAR expression on the cell. **(b)** A total of 1 × 10^11^ AAVs carrying the CD4CAR gene were injected into mice (n=3), and AAV-CD4CAR expression was demonstrated by FACS at the indicated time. Distinct symbols represent different donor PBMCs used for the development of the HuNCG mouse model. All data are shown as the mean ± s.e.m. with error bars on the graph. **(c)** Graphical representation of the biodistribution of AAV-CD4CAR T cells in various tissues (n=3). Bl, blood; Sp, spleen; Li, liver; Lu, lung; It, intestine; Ki, kidney. Cells isolated from the spleen, liver, kidney, blood, and intestine were analyzed by flow cytometry for AAV-CD4CAR expression on CD8+ cells. All data are shown as the mean ± s.e.m. with error bars on the graph. **(d)** Graphical representation of different cytokines detected in the plasma of tumor-bearing ATL NCG-HuPBL mice (n=3), at 2 weeks after AAV injection. Cytokines were detected through FACS by using the BD Cytometric Beads Array Human Th1/Th2 Cytokine Kit. An unpaired two-tailed t-test was used for analysis. ns, no significance; ****P < 0.0001. ***, P < 0.001; **, P < 0.01; *, P < 0.05.

CAR T cells combine features of cell therapy, gene therapy, and immunotherapy. Antitumor characteristics, including transduction efficiency, transgene expression levels, stability or retention, cytokine production, are all key to these processes. Next, we evaluated the stability and biodistribution of AAV-CD4CAR T cells in NCG-HuPBL mice. Our results showed that the AAVs efficiently delivered CD4-CAR DNA into host CD3^+^CD8^+^ T cells, which resulted in the production of enough CD4-CAR T cells, which circulate in the host body for weeks (Figure. 3b). We quantified the biodistribution of AAV-CD4CAR T cells in different organs after injecting AAV-CD4CAR into NCG-HuPBL mice. AAV-CD4CAR T cells were distributed in various organs, with the highest concentrations in the spleen and blood (Figure. 3c). Cytokine production is a hallmark of CAR T cell therapy; therefore, we assessed cytokine production in an ATL NCG-HuPBL mouse model. Our results showed that *in vivo*-generated AAV-CD4CAR cells were robustly functional against tumor cells and were able to produce a series of cytokines (Figure. 3d). Taken together, the findings from these experiments suggest that an AAV encoding a CAR gene can reprogram sufficient immune effector cells to generate *in vivo* CAR T cells. Furthermore, the *in vivo*-generated AAV-CD4CAR T cells can circulate in the host body for weeks and induce antitumor immunological characteristics. These results indicate that the ACG strategy can generate CAR T cells *in vivo*.

### Functional characterization of *in vivo*-generated AAV-CD4CAR T cells

Encouraged by the expression and antitumor immunological characteristics of *in vivo*-generated AAV-CD4CAR T cells, we next decided to compare the functional ability by comparing the CD3^+^CD4^+^ cell depletion and CAR expression of *in vivo*-generated AAV-CD4CAR T cells with those of conventional CAR T cells. We prepared NCG-HuPBL mice transplanted with PBMCs (Figure. 4a) and divided them into four groups: PBS group (untreated control), AAV-CD20CAR group (negative control), AAV-CD4CAR T cell group (positive control), and AAV-CD4CAR group (treatment group). Prior to any treatment, NCG-HuPBL mice were randomly divided into each group with no significant difference in CD3^+^CD4^+^ percentage at day 0 (Supplementary Figure. 3a). We treated the positive control mice with six million T cells infected *in vitro* with AAV-CD4CAR. This amount corresponds to the approved high dose of CAR T cells currently being used in clinics, in which patients are administered 2 × 10^6^ CAR T cells/kg of mean body weight [21]. However, we infused a single dose of 1 × 10^11^ AAV into the treatment group (AAV-CD4CAR) and a negative control group (AAV-CD20CAR). As envisioned, a single dose of AA V (1×10^11^) infusion was able to yield enough AAV-CD4CAR T cells *in vivo*, similar to the number of *ex vivo* transduced cells after infusion (Figure. 3b and 3c). Correspondingly, CD3^+^CD4^+^ cells started decreasing within the first week in the positive control and treatment groups. After two weeks, there was a significant difference in CD3^+^CD4^+^ cell percentage between the positive control and treatment group compared to the negative control and untreated mice (Figure. 4d). However, there was no difference observed between the positive control and the treatment group, suggesting that the targeting potential of the *in vivo*-generated AAV-CD4CAR T cells was similar to that of the conventional CAR T cells. Similarly, no significant difference was observed in percent survival between the positive control mice and the treatment group (Figure. 4e). In summary, the above results show that *in vivo*-generated AAV-CD4CAR T cells are functionally potent and effective against the target in NCG-HuPBL mice and have similar efficiency to that of conventionally manufactured CAR T cells.

**Figure 4.**
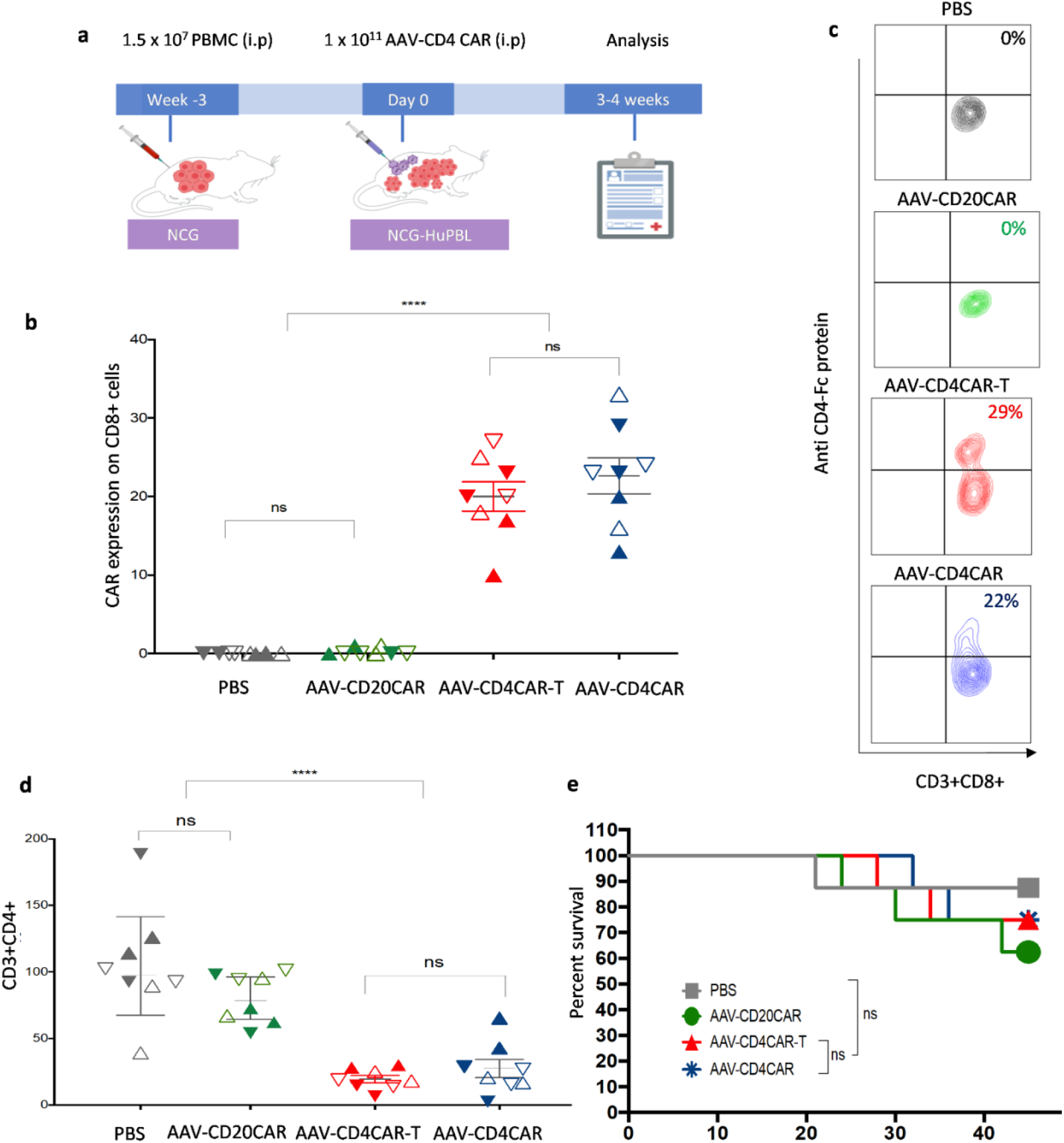
Functional characterization and comparative study of *in vivo*-generated AAV-CD4CAR T cells. **(a)** Schematic representation showing the development of NCG-HuPBL mouse model and experimental design. **(b)** Graphical representation of CAR expression using the CD4-Fc protein in different groups one week after the treatment. The NCG-HuPBL mouse model was developed by using PBMCs from four different healthy donors. Successful NCG-HuPBL mice were randomly divided into four groups, and each group contained eight mice from four different donors. Distinct symbols are used to differentiate the NCG-HuPBL mice engrafted with different donor PBMCs. The treatment regimen was started at day 0 as specified for each group in the main text. Blood was collected from each mouse and evaluated for CAR expression through FACS. Data were analyzed by using two-way ANOVA for multiple comparisons. ****P < 0.0001, ns, No significance. **(c)** Exemplary counter plot showing the CAR expression of a single mouse sample from each group. **(d)** Graphical representation of the normalized CD3+CD4+ percentage documented after 14 days of treatment. Two weeks following the treatment, peripheral blood was collected from each mouse and analyzed for the CD3+CD4+ population through FACS. The CD3+CD4+ cell percentage of each mouse was normalized to 100% at day 0. The CD3+CD4+ percentage at week 2 for each mouse was calculated by the percent difference from the corresponding mice at day 0. Data were analyzed by using two-way ANOVA for multiple comparisons. ****P < 0.0001, ns, No significance. **(e)** Kaplan–Meier curves for percent survival of mice used in (A and B). Percent survival between the groups was analyzed by the log-rank test; P < 0.05 was considered significant.

Next, we also evaluated the relative effects of the ACG approach on blood count, serum chemistry, and mean body. We calculated the bodyweight at different time intervals and collected blood for complete blood count (CBC) and serum chemistry. There was no difference in the mean body weight between the *in vivo*-generated AAV-CD4CAR T group and the untreated group (Supplementary Figure. 3b). Furthermore, serum chemistry and complete blood count values were normal and almost similar to those of the untreated mice (Supplementary Figure. 3c, d). The slight difference in the AAV treated group compared to the PBS group may be due to cytokines production in AAV treated groups. In summary, these results suggesting the safety of the ACG approach for the generation of *in vivo* CAR T cells.

### *In vivo* antitumor activity of ACG

We further assessed whether the *in vivo*-generated AAV-CD4CAR T cells could eradicate the tumor in NCG-HuPBL mice. To establish a tumor-bearing NCG-HuPBL mouse model, we first injected PBMCs into NCG mice and selected successful PBMC-engrafted NCG-HuPBL mice, followed by intraperitoneal injection of luciferase-expressing MT2 ATL cells to develop an ATL NCG-HuPBL mouse model. Mice were randomly divided into two groups: the untreated group (PBS) and the treated group (AAV-CD4CAR). Two days after establishing the ATL NCG-HuPBL mouse model, we injected 2 × 10^11^ AAVs carrying the CD4-CAR gene into the treatment group (Figure. 5a). Interestingly, the ACG strategy started causing tumor reduction at day 10, which became more apparent at day 17 and day 25 (Figure. 5b, c). The mean tumor area of AAV-CD4CAR-treated mice was significantly reduced compared with that of the untreated group (Figure. 5d). Four out of six mice in the treated group presented complete tumor remission. One mouse (M10) showed tumor recurrence when analyzed at day 25 (Figure. 5b). In the untreated group, none of the mice showed any regression in tumor burden, suggesting stable tumor growth over time (Figure. 5b). As CD45^+^CD4^+^ cells are present in both healthy PBMCs and MT2 ATL cells, we next analyzed the CD45^+^CD4^+^ population in mice at the end of the study (day 25) to evaluate the tumor burden at the cellular level. Our results showed that the percentage of CD45^+^CD4^+^ cells in the untreated NCG-HuPBL ATL mice was higher than that in the untreated NCG-HuPBL mice (NCG mice engrafted with PBMCs only, no tumor) (Figure. 5e). This higher percentage of CD45^+^CD4^+^ cells was due to the injected tumor cells that expressed CD4^+^ cells. Interestingly, the FACS results also showed that the CD45^+^CD4^+^ cells were completely depleted in the ACG-treated mice, while no depletion was observed in the untreated mice (Figure. 5f). Furthermore, a significant percentage of CD3^+^CD8^+^ cells showed AAV-CD4CAR expression in the treated group (Figure. 5f). The CD45^+^CD4^+^ depletion and CAR expression in the CD3^+^CD8^+^ population agreed with the tumor burden analyzed by IVIS signal and tumor size in each mouse (Supplementary Figure. 4 a, b). Next, we evaluated the tumor burden at the organelle level. Two mice from each group were euthanized at the end of the study, dissected, and analyzed for IVIS signals (Supplementary Figure. 4 c). Tumors were almost completely eradicated from all organs in AAV-CD4CAR-treated mice, except in one mouse (M10), which showed a small tumor burden in the intestinal region (the same mouse that had tumor recurrence). However, a significant amount of tumor was observed in all the organs in the untreated mice (Figure. 5g). In summary, our results showed that when an AAV carrying the CD4-CAR gene was infused into mice, it could quickly reprogram lymphocytes and generate sufficient CAR cells to induce ATL regression in NCG-HuPBL mice, suggesting that ACG could be a potential strategy for tumor treatment.

**Figure 5.**
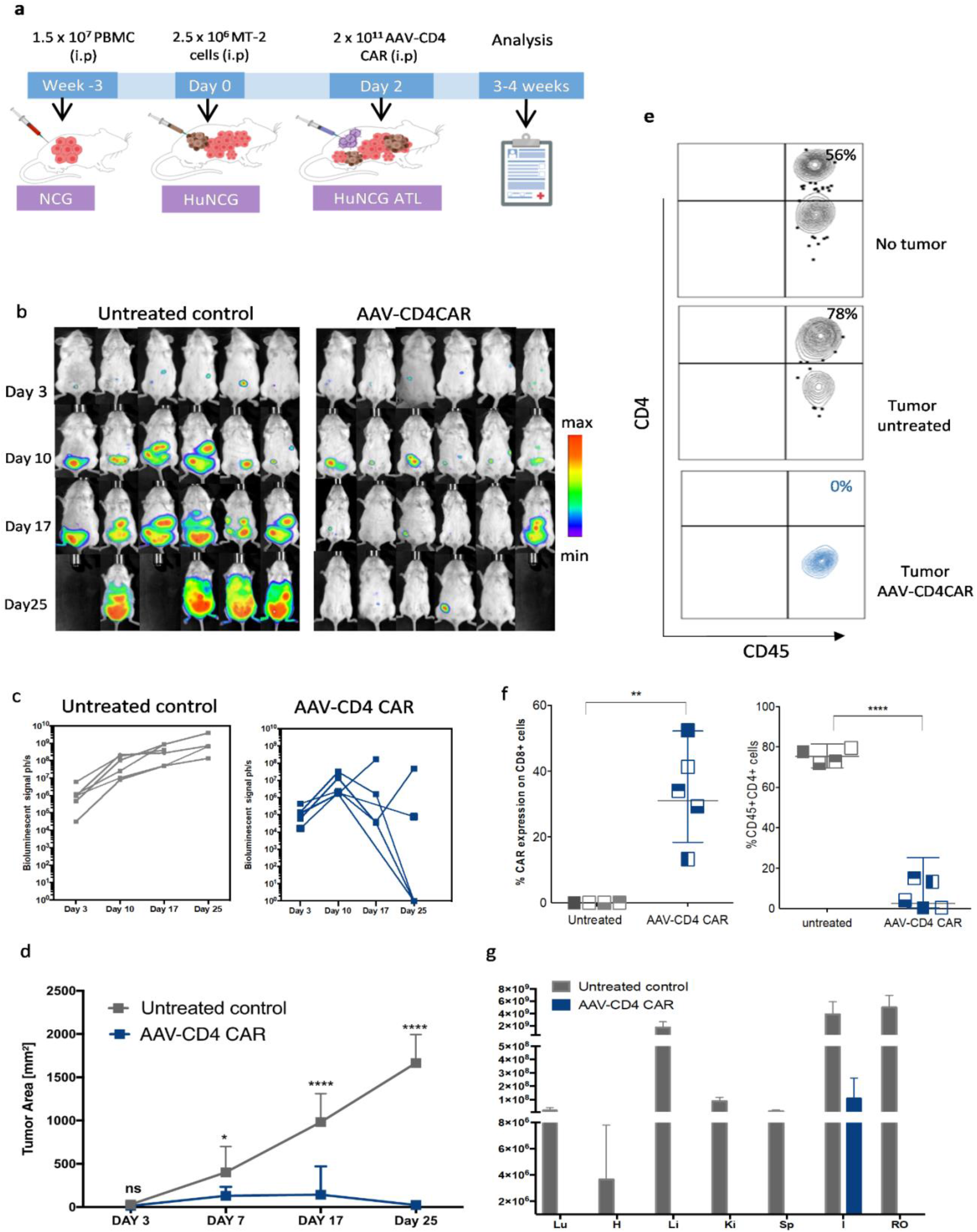
*In vivo*-generated AAV-CD4CAR T cells mediated antitumor activity. **(a)** Schematic representation showing the development of HuNCG ATL mouse model and experimental design. **(b)** Sequential *in vivo* imaging of luciferase-expressing MT2 ATL cells injected into NCG-HuPBL mice. Following AAV or PBS (untreated) injection, mice were imaged the next day, followed by *in vivo* imaging at different t ime points as specified in the figure. Blank images represent dead mice. **(c)** Bioluminescent intensity (p/sec) of mice shown in B. Each line represents one mouse. **(d)** Mean tumor area of AAV-CD4CAR-treated mice and untreated mice. Two-way ANOVA with multiple comparisons was used to analyze the mean tumor area. ns, no significance’ *P<0.05, ****P<0.0001.Data are presented as the mean ± SEM. **(e)** Exemplary counterplot of FACS analysis showing the CD45+CD4+ percentage in NCG-HuPBL mice (no tumor, no treatment), untreated NCG-HuPBL tumor challenge mouse and AAV-CD4CAR-treated NCG-HuPBL tumor challenge mouse at the end of the study. **(f)** The CD45+CD4+ percentage and AAV-CD4 CAR expression of the mice in B. Blood was collected from each mouse at the end of the study a nd analyzed by FACS for the CD45+CD4+ percentage and CAR expression on CD3+CD8+ cells. An unpaired t-test was used to analyze the data. **P<0.001, ****P<0.0001. **(g)** Graphical representation of tumor distribution in different organs. Two mice from each group were selected and sacrificed at the end of the study (Supplemental Figure 4). Organs from dissected mice were imaged immediately after sacrifice. The bioluminescent intensity (pH/s) of each organ was quantified and presented as the mean ± SEM. Lu, Lung; H, Heart; Li, Liver; Ki, Kidneys; Sp, spleen; I, Large and small intestine; RO, Reproductive organs.

## Discussion

AAV has been intensively investigated for gene therapy in preclinical and clinical studies. AAV vectors are among the most suitable tools for *in vivo* gene delivery due to their superior infection efficiency and unique biological and biophysical properties [22]. AAV shows persistent transgene expression and long-term correction of disease phenotype with little or no toxicity in clinical and preclinical studies [23]. In our current study, we used AAV equipped with a synthetic gene to manipulate immune cells. To the best of our knowledge, this is the first report describing that an AAV carrying a CAR gene can reprogram immune cells *in vivo* to generate enough CAR T cells to induce tumor regression. Our current CAR gene-carrying AAV strategy contrasts with the costly and time-consuming method of conventional manufacturing CAR T therapy, which requires primary T cell isolation and transgene introduction and expansion via lentiviral vectors or retroviral vectors *ex vivo*.

We used humanized NCG-HuPBL mice throughout our study, which not only recapitulate ATL but also other antitumor immunological characteristics [24]. In our current study, we chose CD4-targeting CAR transgenes because CD4^+^ leukemia and lymphoma are highly aggressive types of tumors that are extremely refractory to chemotherapy. Initial preclinical studies of CAR T cells for targeting CD4^+^ tumors are encouraging, but to date, no standard care has been developed [25]. Furthermore, CD4+ cells are the major latent reservoir of HIV, posing a challenge for HIV eradication; therapies targeting CD4^+^ cells can also be translated into HIV treatment [26] and treatments for other infectious diseases, including systemic lupus erythematosus, rheumatoid arthritis, multiple sclerosis and psoriasis [27-30]. Previously, our group tested the same targeting antibody (Hu5A8) in humanized N SG-HuPBL mice for HIV treatment [10].

An interesting outcome of our study was that AAVs are capable of producing enough *in vivo* CAR T cells upon a single infusion, and these T cells are potent as *ex vivo*-generated CAR T cells. Furthermore, once produced, they act as a living drug, are distributed throughout the host, and circulate in the host body for weeks, with the ability to recognize and destroy target cells. Conventional lentivirally or retrovirally transduced cells tend to lose their transgenes expression over time [31]. AAV vectors lack engineered lipids and chemical components and are free from viral genes, and thus considered to be safe for gene therapy with limited toxicity [32]. In general, AAV has been reported to be less immunogenic than other viruses and synthetic carriers.

The complete CD4^+^ depletion in NCG-HuPBL mice raises concern for adverse effects in the long term or clinical setting. CD4 helper T cell depletion for long periods leads to “on-target, off-tumor” toxicities and immunodeficiency in patients [33]. Furthermore, the persistence and production of memory AAV-CD4CAR T cells may further extend CD4 cell aplasia in the clinical setting, thereby suggesting the need for a safety switch. One such approach is to eliminate CAR T cells completely from the host. Ma et al. [34] used alemtuzumab (anti-CD52 antibody), which resulted in eliminating CAR T cells within a few hours. Furthermore, such a situation can also be tackled in advance by integrating recently documented multiple advanced logic and control features for CAR T cells [35, 36] into our ACG system.

CAR T cells have emerged as a promising therapy for the treatment of various tumors during the last few years and have obtained approval from the FDA to treat large B cell lymphomas such as non-Hodgkin lymphoma and B cell ALL [1]. Currently, multiple clinical trials and preclinical studies are ongoing to expand the therapeutic applications of CAR T cells. As mentioned earlier, CAR T cell production is a multiplex system involving cell therapy, gene therapy, and immunotherapy. This multiplex system of CAR T cell production limits the therapeutic potential of CAR T cells to a few tumor types and specialized centers around the globe [37]. Several strategies are currently being developed to generate allogenic CAR T cells, including genome editing of T cells [38], *in vivo* engineering of T cells relying on the infusion of nanoparticles [39], and strategies involving lentivirus [40], using alternative effector cells (NK cells, γδ T cells, and macrophages) [41, 42], donor-derived allogeneic CAR T cells [43], and nonalloreactive T cells [44] as well as the split, universal, and programmable CAR system (SUPRA) [35]. All these approaches bear advantages and disadvantages [45]; however, we believe that the AAV-based *in vivo* approach may be simpler to operate in a clinical setting since it transduces dividing and nondividing cells without requiring a strong T cell activation signal. Unlike, Lentivirus and synthetic vector, AAV DNA can persist in episomal form and used host endoplasmic reticulum for uncoating. The episomal form of the AAV DNA is very stable and cannot be degraded by exonucleases [46], thus making it a suitable carrier for manipulating in vivo cells. Additionally, ample evidence from various AAV-based clinical studies has shown the safety of AAVs delivering genes [47].

Our ACG strategy involving *in vivo* generation of CAR T cells has the potential to overcome the abovementioned hurdles and will make CAR T cell treatment easier and affordable, and will transform an intricate, individualized treatment into a broadly applicable product. AAV-DJ subtype for manipulating in vivo immune cells exhibits higher transduction efficiency and broader tropism [11, 48]. Therefore, upon infusion in humanized mice, AAV-CD4 CAR based on AAV-DJ subtype might infect numerous host and target cells. However, the current study aims to give a proof of concept of the ACG method; thus, we evaluate the in vivo generation and properties of key effector cells for CAR therapy, i.e., T cells. Despite our strong proof-of-concept of ACG, there are still many concerns and limitations in our current study, which need to be addressed in the preclinical setting to make these *in vivo*-generated AAV-CAR T cells as effective and safe as *ex vivo*-generated CAR T cells. One major limitation of the current study is the specificity of AAV carrying the CAR gene, and we did not evaluate the CAR expression in hematopoietic cells, tumor cells, and other immune cells. Furthermore, the current study is the lack of having strong data to evaluate the biocompatibility and safety of the ACG approach, and the Hu-PBL NCG may not fully recapitulate the human immune microenvironment. Further studies will be required to design an ACG with ligand coupling or modification in the AAV capsid for cell-specific delivery of the CAR gene. Additionally, a deep insight into its biocompatibility and safety in larger and versatile animal models, and evaluating its therapeutic potential in other tumor models will facilitate clinical translation of this approach. In addition to the in vivo CAR T cells generation, given further optimization, our ACG strategy may lay the foundation for *in vivo* T cell manipulation in immunotherapy beyond cancer and infectious diseases. We believe that the ACG strategy will substantially streamline the manufacturing process of cell-based therapies in the clinical setting and will make genetically engineered cell treatment less expensive and more effective.

## Supporting information

This file includes: Supplementary Methods, Figures S1 to S4, Table S1 and S2, SI References

## Findings

This work was supported by the National Science Foundation of China (NSFC) (No. 81803414, 31970149), The Major Research and Development Project (2018ZX10301406), Jiangsu Province Natural Science Foundation for Young Scholar (grant #BK20170653), Nanjing University-Ningxia University Collaborative Project (grant #2017BN04), Jiangsu Province “Innovative and Entrepreneurial Talent” project and Six Talent Peaks Project of Jiangsu Province.

## Data availability

Plasmids generated in this study will be made available upon reasonable request from the corresponding author. Detailed methods, materials, and troubleshooting strategies are provided as Supplementary Information.

## Ethics

The study and the protocol for this research were approved by the Center for Public Health Research, Medical School, Nanjing University. All experimental animal procedures were approved by the Committee on the Use of Live Animals by the Ethics Committee of Nanjing Drum Tower Hospital. All the authors declare their compliance with publishing ethics.

## Author contribution

All authors contributed to the work fulfilling the criteria adopted from ICMJE. Acquisition of data: WN, WX, BH, SX, ZR, ZL, and YL. Analysis and interpretation of data: WN, WX, SX, and YL. Drafting of the manuscript: WN, WX, and WZ. Critical revision: WX and WZ. Study conception and design: WX and WZ. Financial support: WX and WZ. All authors read and approved the submitted version of the manuscript. Each author has agreed both be personally accountable for the author’s own contributions and to ensure that questions related to the accuracy or integrity of any part of the work, even those in which the author was not personally involved, are appropriately investigated and resolved and that the resolution is documented in the literature.

