## Supplementary material for "AAV-Mediated *In Vivo* CAR Gene Therapy for Targeting Human T Cell Leukemia": This file includes: Supplementary Methods, Figures S1 to S4, Table S1 and S2, SI References

Supplementary Methods

Figures S1 to S4

Table S1 and S2

SI References

#### Supplementary Methods

##### Quantification of AAV by qPCR using SYBR Green technology

We used AAV qPCR to determine the number of genome-containing particles of an AAV prep using SYBR green technology. We treat the purified AAV samples with DNase I to eliminate any contaminating plasmid DNA carried over from the production process (DNase does not penetrate the virion). Removed the extra-viral DNA by digestion with DNase I. Mix 5µl of AAV sample, 5µl of 10 x DNase I buffer, 10µl of DNase I, and 30µl of nuclease-free water. Incubate at 37 ºC for 10 min, 65 ºC for 10 mint to inactivate the enzyme. Dilute DNase-treated AAV samples according to the dilution scheme in the Supplemental Table 1.

| Dilution serious | Volume of the sample (µl) | Volume of nuclease free water (µl) | Dilution factor | Total dilution |
|---|---|---|---|---|
| 1 | 5µl | 90 µl | 10x | 10x |
| 2 | 10µl Dil 1 | 90 µl | 10x | 100x |
| 3 | 10µl Dil 2 | 90 µl | 10x | 1000x |
| 4 | 10µl Dil 3 | 90 µl | 10x | 10000x |

**Supplementary Table 1.** DNase-treated AAV samples for PCR reaction.

Using an AAV-GFP plasmid that contains CMV sequences, prepare 5-fold serial dilutions ranging from 8.7 X $10^{10}$ to 1.1 X $10^{6}$ dsDNA molecules/µl. Count the number of samples (n) and prepare a master mix for an additional 5 samples. In a final volume of 20 µl, mix 2 µl of the template (standard dilution or sample dilution), 0.4 µl CMV-Fwd (10µm), 0.4 µl CMV-Rev (10µm), and 10 µl of the SYBR Green Master Mix and 7.2 µl of nuclease-free water. Each reaction was prepared in triplicate, and no template control (*NTC = master mix + water) was also added.* AAV-Mono-CMV-F: Primer for forward CMV sequence (CCATTGACGTCAATGGGTGGAGT) AAV-Mono-CMV-R: Primer for reverse CMV sequence (GCCAAGTAGGAAAGTCCCATAAGG)

Run the following protocol in your qPCR instrument using SYBR detection:  98°C 3 min / 98°C 15 sec / 58°C 30 sec / read plate/ repeat 39x from step 3 / melt curve. Data analysis was performed using the instrument's software. Determine the physical titer of samples (viral genomes (vg)/mL) based on the standard curve and the sample dilution.

**Western blotting analysis**

HEK-293T cells infected with AAV-GFP or AAV-CD4 CAR were harvested after 48 hours, lysed in 1 mL RIPA buffer with deoxycholate and protease inhibitor cocktail (Roche) for 30 min. Protein supernatant was collected after centrifuging at 13,000 g and 4 °C for 30 min. Protein concentration was determined via the Bradford protein assay (Bio-Rad). Samples containing an equal volume of cells were mixed 1:1 with reducing loading buffer and boiled for 5 min. All lysates were separated on 10% SDS-PAGE gels, transferred to nitrocellulose membrane using the wet cell method, and probed with 1:1000 dilution of mouse anti-human CD3z mAb (Thermo Fisher) and 1:1000 dilution of sheep anti-mouse 800 diluted in PBS-Tween/5% skimmed milk powder. Blots were imaged with an *Odyssey* Imager 600.

**CD4 depletion assay in various donors.**

Blood was obtained from screened healthy volunteers via Blood Bank under informed consent, according to the protocol for research use approved by the medical school of Nanjing University. Donations were fully anonymous to us. PBMCs were purified by Ficoll-Plaque Plus (manufacturer) density centrifugation, as mentioned in the main text in detail. PBMC were recovered from the gradient, washed first with saline, and twice in RPMI 1640 (Gibco-BRL) plus streptomycin-penicillin (sigma) and 10% fetal bovine serum. Viability was assessed by trypan blue exclusion. Washed cells were suspended in RPMI 1640 culture medium (Sigma) containing 10% FBS (Sigma), counted in a hemocytometer, and placed at a level of $2 \times 10^5$ cells per 0.1 mL in flat-bottom 96-well microtiter plates (corning). The next day PBMC were infected by purified AAV (AAV-CD4CAR or AAV-GFP) by adding to each well by gently swirling/mixing in different doses and incubated at 37$^o$ in the presence of 5% $CO_2$. Two days after infecting the PBMC with AAV-CD4 CAR or AAV-GFP, PBMC were phenotyped with anti-Human CD3, anti-human, CD8 and anti-human CD4, as mentioned in detail above. Flow cytometry analysis was performed using a NovoCyte FACS. The percentage of CD4 and CD8 T cells were calculated; the percent changes of CD4 and CD8 in each donor were calculated by normalizing the AAV-GFP percentage to 100.

**Intracellular staining of cytokines**

T cells expressing CAR were co-culture with Jurkat cells in fresh RPMI medium and supplied with IL-2 (300UI/ml) and brefeldin A, and incubated at 37C. After 6 hours, cells were stained for CAR expression. Following washing, cells were fixed and permeabilized by fixation/permeabilization solution (BD). Cells were then stained with anti-Human IFN-γ or anti-Human TNF-α–or anti-Human IL-10 for intracellular staining and incubated at 4C for 30 minutes. Following 30 minutes incubation, cells were washed with BD Perm/Wash buffer and measured on a NovoCyte.

**Generation of stable luciferase-expressing MT2 ATL tumor cell line**

Lentivirus was produced by co-transfection, filtered and titrated as described previously (1, 2). Lentivirus with polybrene (8 μg/ml) was added in MT2 ATL tumor cells and was spin for 3 hours at 32 °C and 200 rpm. The next day above viral transduction was repeated. Cells were grown for 3 days, and the transduction efficiency was evaluated on luminometer with promega Luciferase

Assay System.  Cells were plated in a fresh tissue culture plates to obtained positive cells. Cell clones were grown for ~14 days in the presence of puromycin. Cells were monitored periodically and change fresh media when required. Cells were checked for luciferase activity through a luminometer and IVIS system, and measure its luminescence.  Next, we assessed whether the MT2 ATL tumor cells can grow efficiently *in vivo*. We inject different doses of MT2 ATL cells (0.5 $x10^6$- $5x10^6$) luciferase cell into mice through I.P and S.C. We do a time-course analysis of tumor burden in mice through bio-imaging.

#### Toxicity studies

To evaluate the potential toxicity of AAV in vivo, we divided 15 Hu-NCG mice into 3 groups. In one group, we injected $1 \times 10^{11}$ AAV in each mice. In another group we inject $1 x10^7$ T cells expressing CAR in mice. Whereas, the control group didn’t receive any treatment. Each experimental group comprised of 5 mice. 2 weeks after the AAV injections, mice were anesthetized, and blood was collected by retro-orbital bleed using heparinized microcapillary tubes into ethylenediaminetetraacetic acid-containing microcontainers for determination of the complete blood count (CBC), which included a white blood cell count with differential, a red blood cell count, haemoglobin, haematocrit, and a platelet count.

### Supplementary Figures Legends

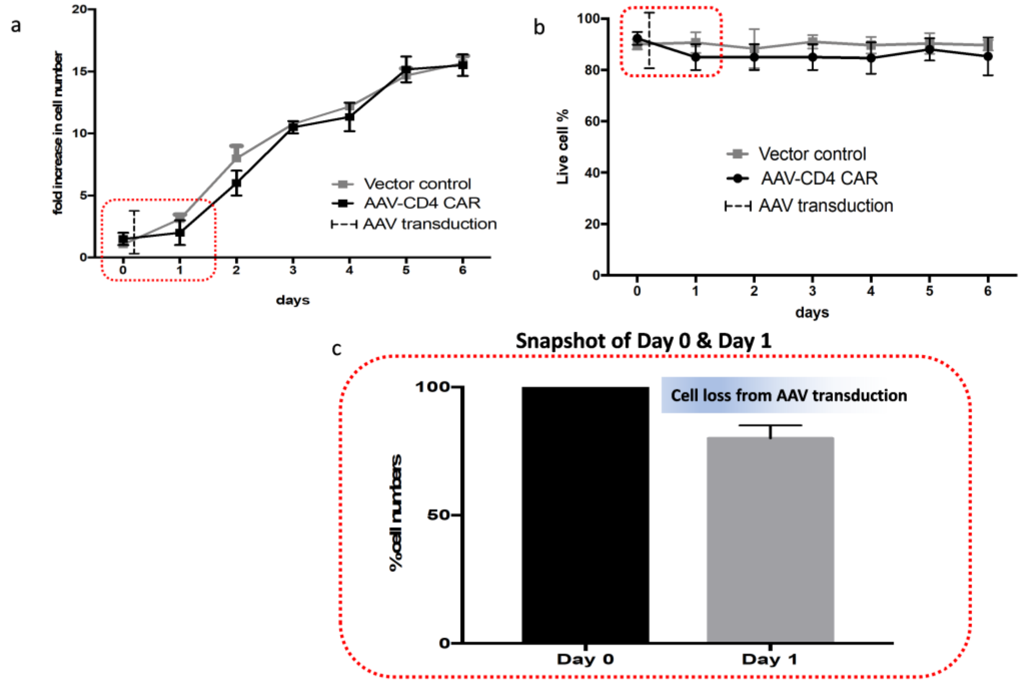

**Supplementary Figure 1. Cell growth and viability testing before and after infection with AAV (cell culture replicates, n = 3). (a)** Time course analysis of cell growth. The total fold increase of cells is shown along with different time points for 7 days. Data are shown as mean ± s.e.m. **(b)** shows the percentage of live cells along a time course. Cell viability was evaluated through trypan blue staining. Statistical analysis showed no significant difference across all time points. Data are shown as mean ± s.e.m. (**c**) Snapshot of Day 0 (before AAV transduction), and Day1 (after AAV transduction). Shows the % cell lost due to AAV transduction. Statistical analysis showed no significant difference in cell loss before and after AAV transduction.

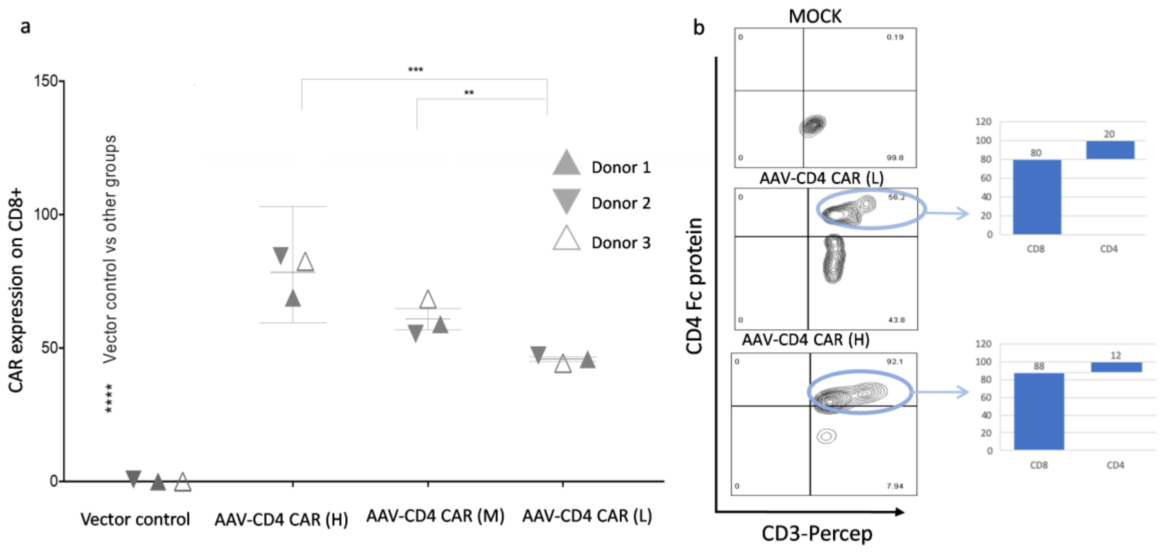

**Supplementary Figure 2. Dose-dependent AAV CD4CAR expression on CD3+CD8+ cell and CD3-CD56+ (NK) cells. (a)** PBMC were activated with IL-2. Cells were infected with different doses AAV-CD4CAR, (AAV-CD4CAR (L) is defined as MOI=1x$10^4$/Cell, AAV-CD4CAR (M) as 0.5 x $10^5$/cell and AAV-CD4CAR (H) as MOI= 1x$10^5$/Cell). Where one set of cells was left uninfected and served as vector control. 48h post-infection cells were stained with CD4-Fc protein, anti-human CD3, and anti-human CD8 antibody, followed by secondary antibody staining with AlexaFlour488 goat anti-rabbit IgG(H+L) for the CAR Expression. A two-way ANOVA multiple comparison were used to calculate the significant difference. **P < 0.001, ***P < 0.001. Different shape triangles represent different donors. **(b)** PMBCs were infected as mentioned above, 48h post-infection cells were stained with CD4-Fc protein, anti-human CD3 and, anti-human CD4 antibody and anti-human CD8 antibody, followed by secondary antibody staining with AlexaFlour488 goat anti-rabbit IgG(H+L) for the CAR expression.

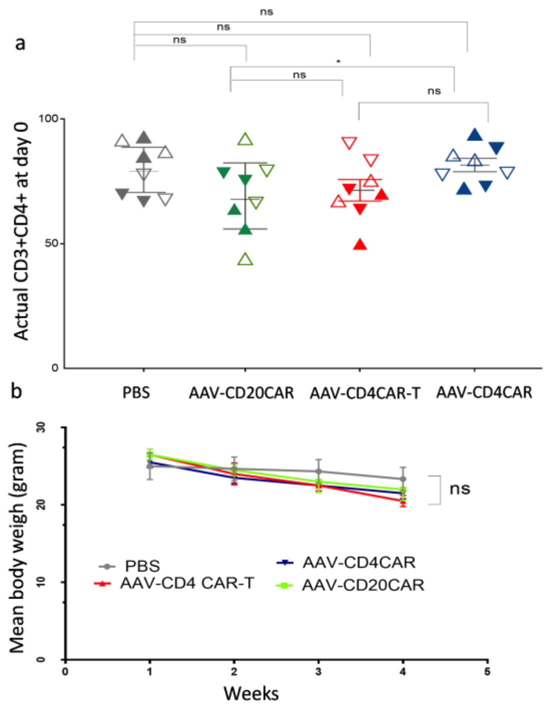

c

Complete Blood count (CBC)

| | PBS | AAV-CD20 CAR | AAV-CD4 CAR-T | AAV-CD4 CAR |
|---|---|---|---|---|
| **WBCs** (10^9/L) | 3.8±0.8 | 3.6±0.9 | 4.1±1 | 3.6±0.9 |
| **RBCs** 10^12/L | 1.91±0.5 | 2.12±0.5 | 2.31±0.9 | 2.12±0.5 |
| **Hb** g/L | 56±12 | 57±4 | 45±3 | 57±4 |
| **Hematocrit** % | 15.3±2 | 21.9±6 | 25.6±8 | 21.9±6 |
| **platelet** 10^9/L | 216±21 | 201±14 | 188±21 | 201±14 |
| **MCV**(fl) | 80.1 ±7 | 75±8 | 59.4±14 | 75±8 |

d

Clinical Blood chemistry

| | PBS | AAV-CD20 CAR | AAV-CD4 CAR-T | AAV-CD4 CAR |
|---|---|---|---|---|
| ALT (U/L) | 11±5 | 29±12 | 23±8 | 19±5 |
| LDH (U/L) | 300±25 | 450±29 | 334±32 | 312±35 |
| AST (U/L) | 56±12 | 57±4 | 45±3 | 57±4 |
| Cr (mmol/L) | 15.3±2 | 21.9±6 | 25.6±8 | 21.9±6 |
| CK (U/L) | 6±2 | 8±2 | 18±3 | 13±5 |
| BUN (mmol/L) | 3 ±1 | 5±3 | 8.5±1.5 | 7.5±0.8 |

**Supplementary Figure 3. Percentage of CD3+CD4+ on day 0 and Safety characterization of AAV treatment.**

**(a)** The percentage of CD3+CD4+ at day 0 for *in vivo* CD3+ CD4+ depletion (shown in Figure 4) in different groups before starting any treatment regimen. Data were analyzed using two-way ANOVA for multiple comparisons. *P < 0.05, ns, No significance. **(b)** Time-course analysis of mean body weight. Data are shown as mean ± s.e.m at a specific time point. Two-way ANOVA with multiple-comparisons test was used to assess significance (uncorrected Fisher's LSD). **(c)** Serum blood chemistry**.** Data are shown as mean ± s.e.m. **(d)** Blood count. Data are shown as mean ± s.e.m. There were no differences in the blood counts and serum blood chemistry of AAV treated groups and the PBS injected group.

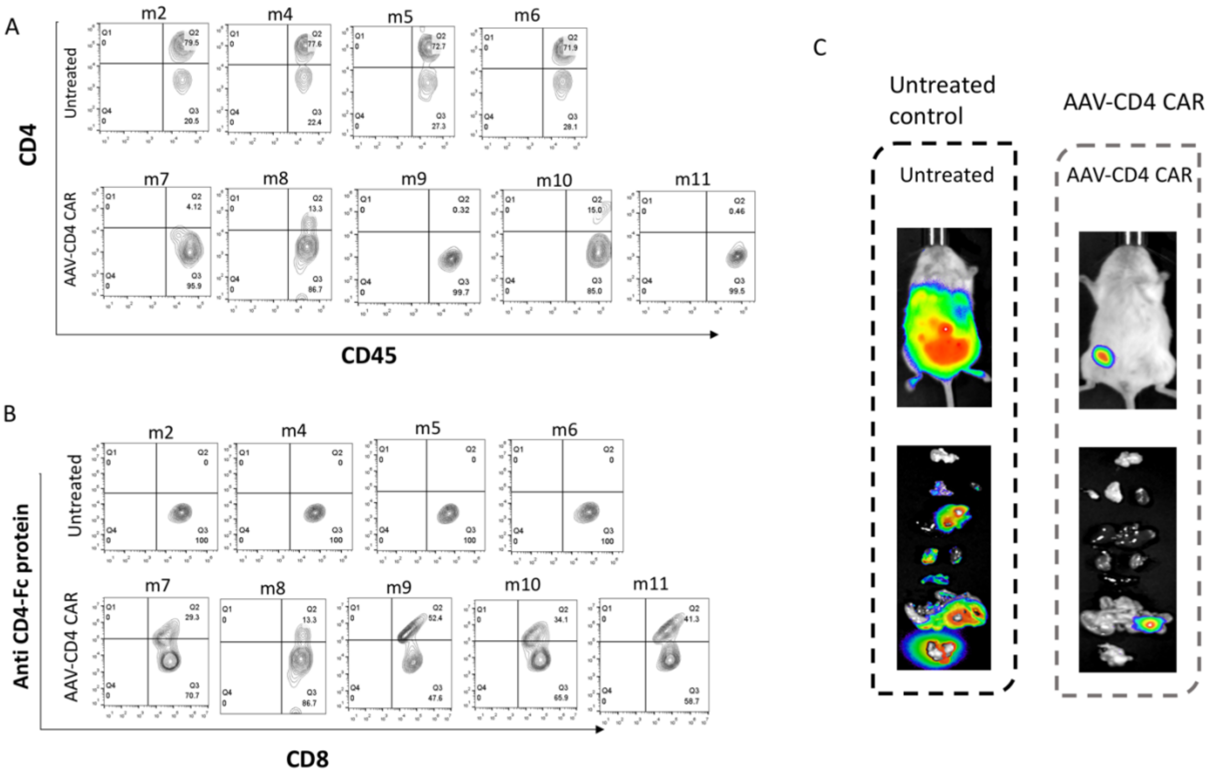

**Supplementary Figure 4. FACS results showing the CD45+CD4+ cells and CAR expression on CD3+CD8+ cells, and representative images of the tumor burden in the mouse at the end of study (a)** FACS result showing the CD45+CD4+ cells from each mouse at the end of the study. M1 represents the first mouse of the untreated group (from left) and then follows this order in right direction. Graphical representation and statistical analysis are shown in Fig. 5f. (**b)** AAV-CD4CAR expression on CD3+CD8+ cells at the end of the study. Blood was collected and stained for CD4-rFc protein (for CAR expression) and other required antibodies. Graphical representation and statistical analysis are shown in Fig. 5g. (m1, m3, and m12 were died before final analysis, and are not included in figure S4 a, b) **(c)**Two mice from each group were selected and sacrifice at the end of the study. Organs from dissected mice were imaged immediately after euthanized. Graphical representation and statistical analysis are shown in Fig. 5g.

**Supplementary Table 2.** List of materials

| Reagent or resource | Source | Identifier |
|---|---|---|
| **Cell lines** | | |
| AAV293 | Agilent | 240073 |
| HEK293 | ATCC | CRL-11268 |
| HEK293F | Thermo Scientific | R79007 |
| Jurkat | ATCC | TIB-152 |
| MT2 ATL | Sigma | 08081401 |
| MT2 ATL-Luc | This paper | N/A |
| PBMC | This paper | N/A |
| **Plasmids** | | |
| pAAV-DJ | This paper | N/A |
| AAV-pHelp | This paper | N/A |
| AAV-CD4CAR | This paper | N/A |
| AAV-CD20CAR | This paper | N/A |
| AAV-GFP | This paper | N/A |
| **Antibodies** | | |
| FITC Mouse anti-Human CD3 clone- | Biolegend | CAT # 300440<br>Clone# UCHT1 |
| Pacific Blue Mouse anti-Human CD3 clone- | Biolegend | CAT # 300440<br>Clone# UCHT1 |
| PerCp Mouse anti-Human CD3 clone- | Biolegend | CAT # 300326<br>Clone# HIT3a |
| APC Mouse anti-Human CD45 clone- | Biolegend | CAT # 304037<br>Clone# H130 |
| PE Mouse anti-Human CD8 clone- | Biolegend | CAT # 344706<br>Clone# SK1 |
| PeCy7 Mouse anti-Human CD4 clone- | Biolegend | CAT# 317414<br>Clone# OKT4 |

| | | |
|---|---|---|
| Anti-Human CD45-APC | Biolegend | CAT# 304037<br>Clone# H130 |
| Anti-Human CD56 Pacific blue | Biolegend | CAT# 362519<br>Clone# 5.1H11 |
| Anti-Human CD45RO Brilliant Violet 421™ | Biolegend | CAT# 304223<br>Clone# UCHL1 |
| Anti-Human CD62L APC/Cy7 | Biolegend | CAT# 304813<br>Clone#DREG-56 |
| AlexaFlour488 goat anti-Rabbit igG(H+L) | Invitrogen by thermosscientific | Ref# A11034 |
| **Experimental Models: Organisms/Strains** | | |
| **NCG** | Genetech Pharma | N/A |
| **Commercial Assays** | | |
| XenoLight D-Luciferin - K+ Salt Bioluminescent Substrate | PerkinElmer | Cat#122799 |
| CBA Human Th1/Th2/Th17 cytokine kit | BD | Cat# 560484 |
| **Chemicals, Peptides, and Recombinant Proteins** | | |
| Phosphate-Buffered Saline | Thermo Scientific | Cat#AM9625 |
| PEG 8000 | Sigma | CAS No: 25322-68-3 |
| PEI | Sigma | Cat#408727 |
| SYBR green | Novozymes | Product No: Q511-02 |
| **Software and Algorithms** | | |
| Graph Pad Prism 7 | Graph Pad | https://www.graphpad.com/scientific-software/prism/ |
| https://biorender.com/ | Biorender | biorender.com/ |

| | | |
|---|---|---|
| FlowJo V10 | TreeStar | https://www.flowjo.com/ |
| Indigo | Berthold Technologies | https://www.berthold.com/en/bioanalytic/products/in-vivo-imaging-systems/nightowl-lb983/ |
| ImageJ | Wayne Rasband | https://imagej.nih.gov/ij/ |
| **Others** | | |
| DMEM | Hyclone | SH30243.01 |
| RPMI-1640 | Hyclone | SH30027.FS |
| OPTI-MEM | Gibco | 31985070 |
| Puromycin | Thermo Scientific | Cat# A1113803 |
| Fetal bovin serum | Excell Bio | FSP500 |
| Phosphate buffer saline | Invitrogen | C20012500BT |
| Brefeldin A | Cayman | 11861 |
| **Instruments** | | |
| BD FACSAria | Biosciences | https://www.bdbiosciences.com/en-us |
| *NovoCyte* | ACEA Biosciences, Inc. | https://www.aceabio.com/ |
| *qPCR* | Applied Biosciences | https://www.thermofisher.com/pk/en/home/life-science/pcr/real-time-pcr/real-time-pcr-instruments.html |
| IVIS Spectrum *In Vivo Imaging System* | PerkinElmer | https://www.perkinelmer.com/product/ivis-instrument-spectrum-120v-andor-c-124262 |
| NightOWL II LB 983 | Berthold Technologies | https://www.berthold.com/en/bioanalytic/products/in-vivo-imaging-systems/nightowl-lb983/ |
